# Photomapping electrically coupled networks of the mature thalamus and cortex

**DOI:** 10.1101/2025.03.06.641890

**Authors:** Mitchell J. Vaughn, David S. Uygun, Radhika Basheer, Kevin J. Bender, Julie S. Haas

## Abstract

Electrical synapses are expressed ubiquitously across the brain and are crucial components of active neural circuitry and connectomes. Identification of coupled networks in living tissue is limited by technical demands of multiplexed recordings, and no dyes, fluorescent reporters, or genetic labels can currently fill the gap. We introduce a novel method of identifying and quantifying electrical synapses, opto-δL, that combines focal photostimulation of soma-targeted opsins with a spike timing-based computation for the strength of electrical synapses to rapidly measure and map electrically coupled networks *in vitro*. We leverage opto-δL to show that coupled networks of the mature thalamic reticular nucleus extend as far as 100 μm, synapse promiscuously across genetic subtypes of neurons, and couple 1-4 neighboring neurons to each recorded hub cell. We also demonstrate application of opto-δL to cortical networks. These results highlight the broad potential of opto-δL to interrogate the identity and roles of electrical synapses in circuitry, behavior, and cognition.

**Teaser:** Electrically coupled neural networks are newly unmasked by functional photomapping.

## Introduction

Chemical and electrical synapses, the twins of the foundational Golgi-Cajal debate, together compose the basis of interneuronal communication across the brain. Neurotransmitter-based chemical synapses underlie communication that is primarily unidirectional and stereotyped in timecourses of activation and decay. In contrast, electrical synapses formed by gap junctions (GJs) relay neuronal signals of any size, polarity, and timecourse bidirectionally between coupled cells, and they do so without the requirement or cost of a spike for signal transduction. Most mature mammalian neuronal GJs comprise paired hexameric channels of connexin36 (Cx36) (*1–3*), which is expressed widely in GABAergic neurons throughout the brain (*4*). While chemical synapses and the neuronal networks they form have been described in exquisite detail, descriptions of electrical synaptic transmission and the functions of coupled networks have been much more limited.

Electrical synapses are increasingly appreciated for the diverse roles they play throughout the brain. They are most widely characterized as mediating synchronization of oscillatory activity in neuronal circuits (*5–15*). Beyond synchronization, electrical synapses directly mediate feedforward and lateral excitation, inhibition, reduce signal to noise ratio, sharpen coincidence detection, and facilitate object discrimination (*16–32*). GJs create uniquely offset receptive fields in retinal ganglion cells, where they also contribute to tuning responses, enhanced signal saliency, sensitivity, and enhanced coding of visual features (*16, 33–36*).

Our understanding of the roles electrical synapses and coupled networks play in the central nervous system is acutely limited by the tools currently available. The gold standard for measuring electrical synapses is through paired recordings, in which current is injected into one neuron and a hyperpolarizing deflection can be measured in a coupled neuron (*37*); the ratio of deflections indicates synapse strength. While this method is the most direct, it is time-intensive, low-throughput, and limited to single synapses *in vitro*. Only one experiment has successfully recorded from a small sample of coupled neurons, close to the brain surface, in rodents *in vivo* (*38*). Alternatively, networks can be distinguished by dye-coupling methods (*39–46*), but these do not differentiate between primary, secondary, or tertiary connections, fail to quantify the strength of synapses, and are limited to fixed tissue. Updated variations of dye-coupling have used exogenously applied molecules that diffuse across a GJ, rendered more specific by limiting entry (*47*) or fluorescence (*48*) to genetically labeled cells. Due to these limitations, electrical synapses and networks remain neglected relative to their chemical counterparts in our evolving understanding of the mammalian brain. Modern connectomes (*49*) have almost entirely neglected electrical synapses, with the exception of the retinal connectome (*50*).

The thalamic reticular nucleus (TRN) is a nucleus of GABAergic neurons (*51*) that provide the majority of inhibition to sensory thalamic relay nuclei and are hypothesized to focus the searchlight of attention upon the sensory surround (*52*). The TRN is composed of distinct neuronal subtypes with spatially and genetically distinct expression (*53–55*), and our previous paired recordings indicated that homocellular and heterocellular electrical synapses (coupling matched and disparate cell subtypes) of the TRN (*56*) may provide the only thalamic link between distinct but parallel thalamocortical relay channels (*53–55, 57*). Because visualization of TRN neurons is impeded by myelination of thalamocortical fibers after eye opening, all paired-patch recordings characterizing electrical synapses in TRN have been limited to juvenile tissue (*12, 56, 58–64*). While Cx36 expression persists into adulthood (*65*) and spikelets have been reported in adults (*66*), the possibility remains that the contributions of GJ networks, and thereby any interconnectivity within TRN, to thalamocortical relay are limited to developmental stages.

Here, we leverage focal photostimulation of a soma-targeted opsin to detect and measure electrical synapses via changes in spike timing, which enabled us to map electrically coupled networks in mature TRN and cortex. Our maps of electrically coupled networks in mature TRN demonstrate that coupled networks comprise both heterocellular and homocellular electrical synapses that range from 1 to 4 synapses per hub neuron and extend as far as 100 μm of distance. In addition to providing a new and efficient toolset for measuring and mapping electrical synapses and networks across the living brain, our results provide new insight into the neural circuits that underpin attentional focus on the sensory surround.

## Results

Precision of neuronal spike timing is a tightly controlled fundamental process (*67*) that is crucial for fidelity of information processing and is often a parameter of interest in investigations of neural coding (*68–72*). Electrical synapses are important contributors in determining spike times (*30, 31, 60*); a quiescent neuron within a GJ-coupled network delays its neighbors’ spikes by shunting current, while an active neuron shortens latencies in neighbors by sending excitatory currents across GJs. Activity in a coupled neighbor can shift spike times of a neuron by tens of ms (*60*). We previously measured changes in spike latencies due to electrical synapses in a cohort of coupled juvenile TRN neurons (*73*) by comparing the means of a set of spike latencies from input in two cases: 1) rheobase stimulation in one neuron alone 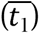 and 2) rheobase stimulation with simultaneous activation of a coupled neighboring neuron 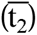. We calculated the percentage change in latency of evoked spikes, □L, as:

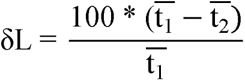

This quantity is well-correlated with traditional measures of the strength of electrical synapses (*73*). Here, we combined measurements of δL from a single patched hub neuron with individual focal photostimulations of nearby neurons expressing a soma-targeted opsin, st-ChroME (*74*), thereby reducing the requirement for recording an electrical synapse to one patched neuron and simultaneously widely expanding the number of potential synapses tested. We refer to this method as opto-δL.

We first validated opto-□L in a patched pair of neurons from juvenile tissue *in vitro* following injection at P0 of AAV9-CAG-DIO-ChroME-st-P2A-H2B-mRuby; both neurons in this pair expressed st-ChroME. We measured the coupling coefficient (cc) for the synapse between these neurons using the traditional small (50-100 pA) hyperpolarizing current injections to one of the neurons (Fig. 1A); cc is the ratio of voltage deflections in both neurons in response to current injection into one of the neurons. This pair had a mean cc of 0.075, close to the average value for coupling between pairs of TRN neurons (*75*). Next, we used a digital mirror device (Mightex Polygon) to deliver 1-photon focal (34 - 50 µm^2^) 560 nm photostimulation to cell 2 of this pair, interleaved during trials of rheobase-driven spiking in cell 1, the patched neuron. We used a light intensity minimally sufficient to drive spiking when focused directly on the soma of cell 1 (see Methods; Supp. Fig. 1). Because both 1-photon and 2-photon photostimulation scatter light in live tissue (*76–79*), we also measured spike times in cell 1 of this patched pair when photostimulating a blank spot that was equally distant from cell 1, as a control for scatter-induced photoexcitation of cell 1. We computed □L from the latencies of spiking in cell 1 elicited during a set of 10 trials of rheobase stimulation, and 10 trials of rheobase with focal photostimulation of cell 2. Driven by rheobase current (100 pA) alone, cell 1 spiked 176 ± 3 ms after stimulus onset (Fig. 1B, dark blue traces). During focal photoexcitation of cell 2, cell 1 spiked with a latency of 99 ± 0.3 ms (Fig. 1B, lighter blue traces), and we computed initial □L for the synapse from the two latencies: □ L = 43.4 ± 0.5%. During photoexcitation of the equidistant blank spot, cell 1 spiked with latency of 148 ± 3 ms (Fig. 1C, light blue traces) and produced □L of 16.3 ± 2.2%. Because □L was significantly larger (by 27.1 ± 1.6%; p < 0.01, Wilcoxon signed-rank test) for photostimulation of cell 2 compared to the blank spot, we conclude that cell 2 was coupled to cell 1, with final corrected □L here taken as 27.1 ± 1.6%. As this example demonstrates, it is critical that all opto-□L measurements are normalized by scatter-control measurements, and all opto-□L values reported here are normalized to controls (see Methods). The values of cc and opto-□L from this example pair were consistent with previous measurements of cc and □L in which both quantities were measured by direct current injections in juvenile tissue (Fig. 1D) (*73*).

**Fig. 1.**
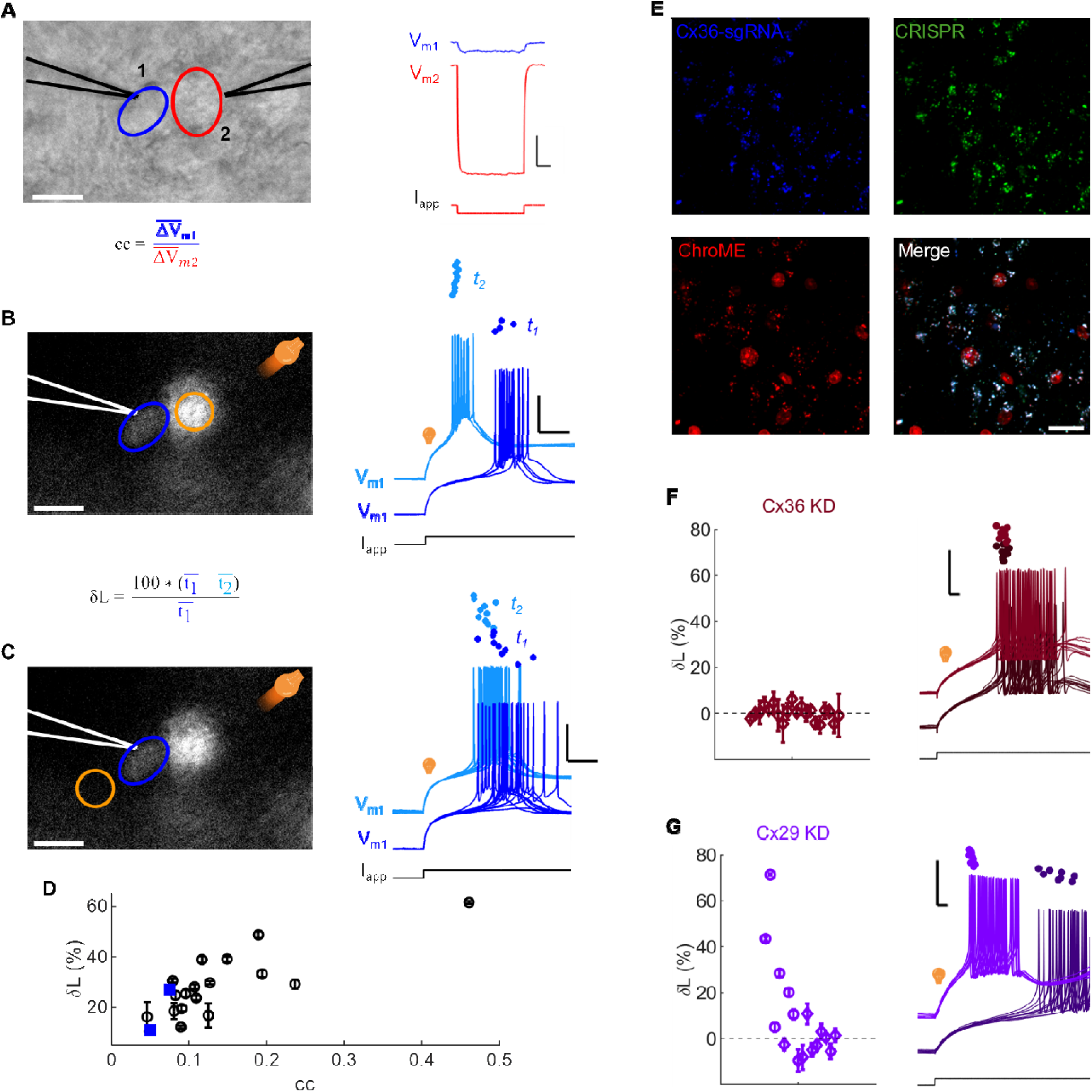
Electrical synapses measured by paired patching and by opto-δL. **A.** Live IR image of a patched pair of TRN juvenile neurons *in vitro*. Right: voltage responses (V_m_) from both neurons during current injection (I_app_) into cell 2. The coupling coefficient (cc) is the ratio of voltage deflections in both cells; cc = 0.075. Scale bars 100 ms, 2 mV. Both cells were maintained at V_m_ = -70 mV by constant current application of -83.3 or -57.5 pA. R_in_ = 82.8 and 88.4 MΩ. **B.** Live fluorescence image of the same pair from **A;** st-ChroME-mRuby was expressed in both neurons. Orange: area of focal photostimulation of cell 2. Scale bar: 12.5 μm Right: Spiking in neuron 1 for 100 pA applied current alone (dark blue), and during simultaneous focal photostimulation of neuron 2 (light blue). Dots mark the times of the first spike in each interleaved trial. δL was initially calculated from the difference in mean spike latencies with () and without () photostimulation; δL = 43.4 ± 0.5%. **C.** Orange oval marks the area of focal photostimulation of a blank spot that is equidistant from neuron 1. Right: Spiking in neuron 1 for 110 pA applied current alone (dark blue), and with simultaneous photostimulation of the blank spot (lighter blue) from interleaved trials. Dots mark the times of the first spikes in each trial. Scale bars 50 ms, 20 mV. For the blank spot, δL = 16.3 ± 2.2%. Differences between δL for trials in **B** and **C** were significant (p < 0.01, Wilcoxon signed-rank test). Corrected δL for this pair: 27.1 ± 1.6%, p < 0.01. **D.** Open circles: cc and δL, both measured by current injections in a cohort of paired recordings (adapted from Haas (2015)). Blue squares: cc and opto-δL from two ChroME-expressing juvenile pairs, including the pair in **A-C**. **E.** Maximum intensity projections of TRN cells from CRISPR knockdown of Cx36 (Cx36 KD). Blue: Cx36-SgRNA-BFP, Green: PV-Cre-Cas9/GFP, Red: ChroME-mRuby. Scale bar: 20 µm. **F.** Left: Measurements of δL (from 20 possible connections to 3 patched neurons from 3 slices) in Cx36 KD. No δL measurements were significant (mean □L = 0.2 ± 0.7%). Right: Example spiking responses used to measure □L in Cx36 KD tissue. Black traces are responses during control trials and colored traces are responses during focal target cell photostimulation. Scale bars: 5 ms, 20 mV. **G.** Left: measurements of δL (from 16 possible connections to 2 patched neurons from 2 slices) from CRISPR knockdown of Cx29. Circles: significant δL values (mean □L = 29.8 ± 10.0%); diamonds: δL not significant (mean □L = -1.7 ± 1.9%) Right: Example spiking responses used to measure □L in Cx29 KD tissue. Black traces are responses during control trials and colored traces are responses during focal target cell photostimulation. Scale bars 5 ms, 20 mV.

We further validated opto-δL using PV-Cre-Cas9 mice injected with AAV5-SgRNA-Cx36-BFP to knock down (KD) Cx36, which expresses in neurons (Fig. 1E). As a control for non-specific genome abscission, we used AAV5-SgRNA-Cx29-BFP to knock down Cx29, which only expresses in glia, where Cas9 is not expressed. In 3 slices prepared from 3 mice, we detected no electrical synapses among 3 patched hub neurons and 20 neighboring neurons using opto-δL in the Cx36 KD (Fig. 1F; mean □L = 0.2 ± 0.7%, p > 0.05, Wilcoxon signed-rank test). In contrast, we detected 6 electrical synapses by opto-δL from 2 patched hub neurons and 16 neighboring neurons in 2 slices from 2 Cx29 KD mice (Fig. 1G; mean □L for all synapses = 10.1 ± 5.4%; mean □L for 6 coupled synapses = 29.8 ± 10.0%, p < 0.05, Wilcoxon signed-rank test). Collectively, these measurements validate the use of a single electrode combined with focal photostimulation of soma-targeted opsins to identify and characterize electrical synapses.

Further, this validation provided a foundation for using opto-□L to identify and measure electrical synapses and networks in mature TRN tissue, where spikelets in depolarized neurons provide evidence of electrical synapses (*66*) (Fig. 2) but heavy myelination precludes paired recordings.

**Fig. 2.**
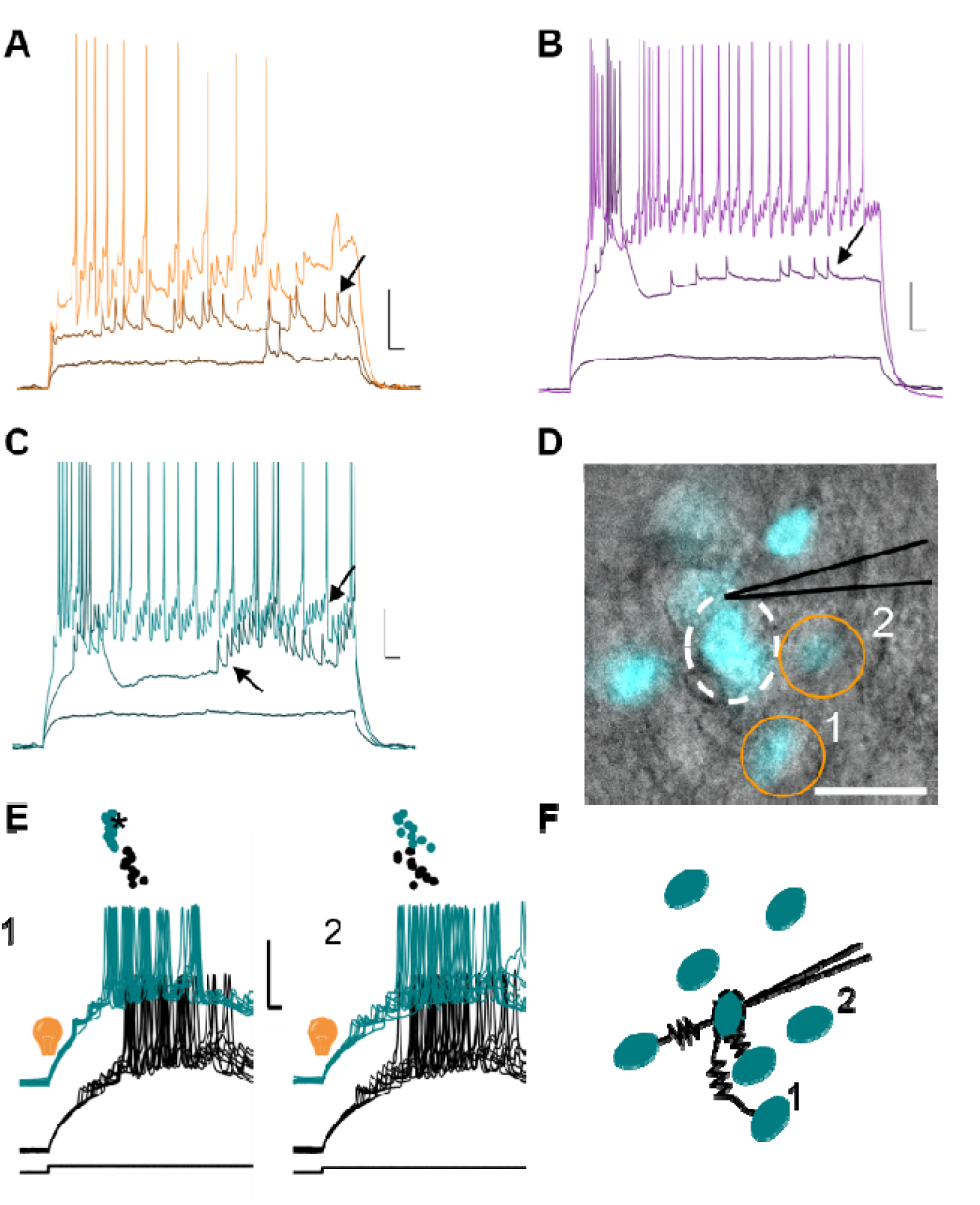
Evidence of electrical synapses in adult TRN. **A – C.** In three different TRN cells from adult tissue, spikelets (arrows) appeared upon depolarizations (by 50, 100 and 150 pA from V_m_ = -70 mV) that presumably excited an unrecorded coupled neuron. Scale bars 10 mV, 25 ms. **D.** Neighborhood of the patched neuron in C (composite IR and live fluorescence image, stChroME-mRuby pseudocolored). Scale bar 25 μm. **E**. Traces from opto-δL measurements of neurons 1 and 2 from **D**, as described **in** Fig. 1B**-C**. In this cluster, opto-δL distinguishes between neuron 2 (δL= 4.9 ± 3.6% Y, p = 0.13) that lacks an electrical synapse to the hub neuron, and equidistant neuron 1 (δL = 25.0 ± 1.2%, p < 0.01) that does have an electrical synapse to the hub neuron). Scale bars 20 mV, 5 ms. **F.** Putative coupled network determined from separate focal photostimulations of each neuron in **D**.

Genetically and anatomically distinct sets of neurons form distinct sets of relay pathways through the thalamocortical system (*53–55, 57*); as GABAergic synapses within TRN do not appear in adulthood (*80*), electrical synapses within the TRN may be the only thalamic link between these information channels. We previously used paired recordings in juvenile tissue to test whether TRN electrical synapses interconnect neurons of distinct cell types (*56*). Those experiments necessitated the use of crossed Cre reporter mice to label neurons for juvenile recordings, presenting an additional possible confound of mislabeling neurons due to Cre off-target recombination (*81*). Therefore, we used opto-□L to identify, measure and characterize homocellular and heterocellular electrically coupled networks in mature TRN *in vitro*, at least 2 weeks after injection of AAV9-CAG-DIO-ChroME-st-P2A-H2B-mRuby into the TRN of SOM-Cre and PV-Cre mice at age P28 – P35. To map homocellular electrical synapses and networks, we patched ChroME-expressing mRuby^+^ hub neurons (SOM^+^) in 36 slices from 34 SOM-Cre mice and measured □L induced by focally photostimulating other nearby mRuby^+^ neurons (Fig. 3A-C; Supp Fig. 2). To map heterocellular electrical synapses and networks, we patched mRuby^-^neurons (SOM^-^) in 13 slices from 12 SOM-Cre mice and measured □L induced by focally photostimulating nearby mRuby^+^ neurons (Fig. 3D-F; Supp Fig. 2). We also measured homocellular networks amongst PV^+^ neurons (Fig. 3G-I; Supp Fig. 2). To compensate for excitation from scatter in these experiments, we developed correction procedures for each hub neuron (Supp. Fig. 3; see Methods); scatter effects depend on factors including light intensity, opsin expression, and input resistance, as well as distance. Depolarization from scatter diminishes to insignificant levels by 50 - 100 μm from the photostimulated area (Supp. Fig. 4). We note that mapping heterocellular networks has the advantage of targeting opsin-lacking neurons for recording, thereby averting the need for compensation in those experiments.

**Fig. 3.**
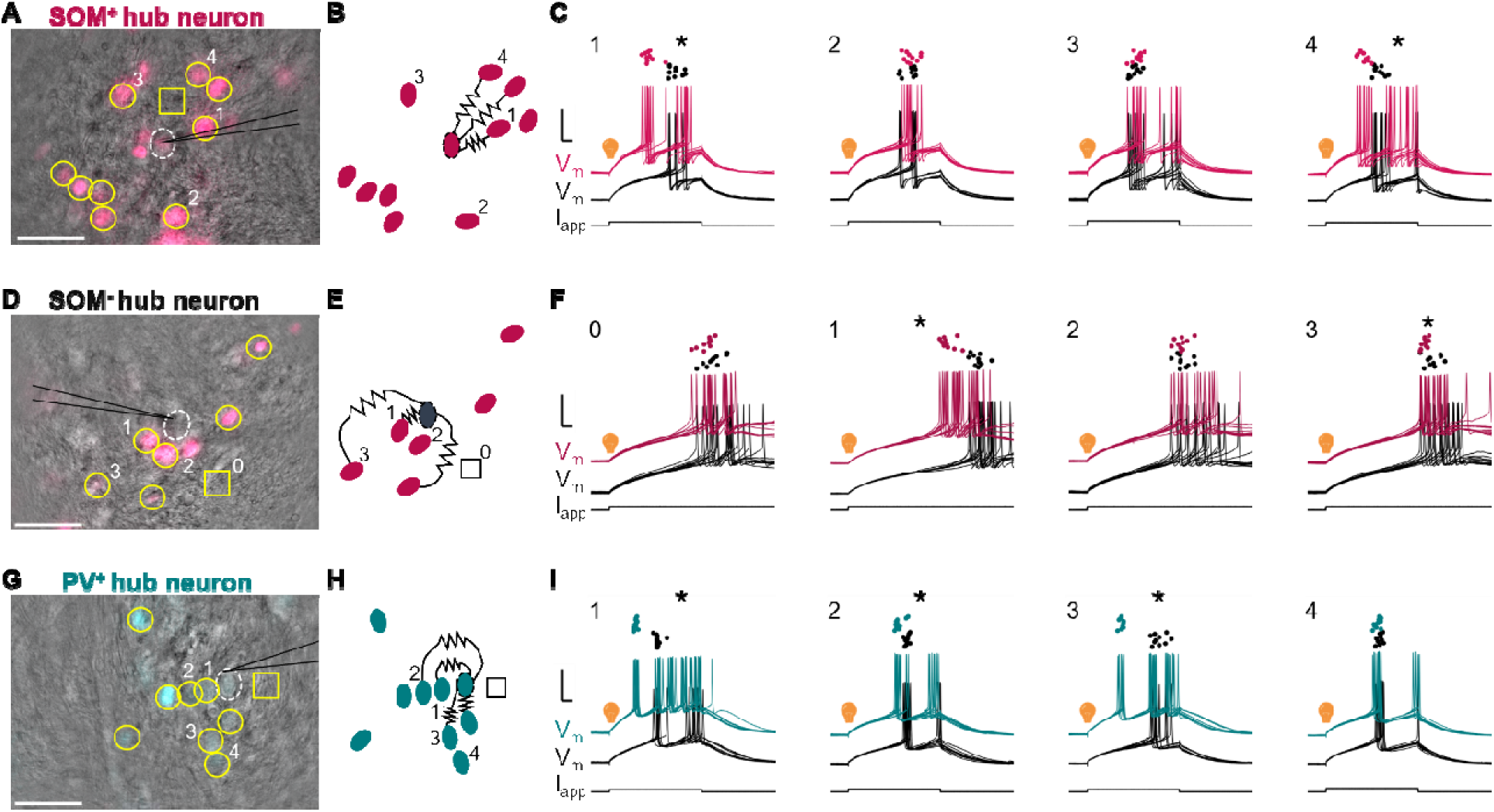
Maps of coupled networks between TRN neurons. **A.** Overlay of IR and fluorescence images from live recording of st-ChroME-mRuby-expressing SOM^+^ TRN neurons *in vitro*; fluorescence is pseudocolored magenta. The patched neuron is circled with white dashes. Cells tested by opto-□L are circled in yellow. One empty control area is marked by a square. Scale bar: 50 μm. **B.** Diagram of the coupled network embedded within all tested neurons from **A**. **C.** Spiking responses from opto-□L measurements from numbered cells in **A**. Dots mark the times of first spikes in the patched neuron from each trial. Black traces are responses of the patched neuron during rheobase-only trials and colored traces are responses during added focal photostimulation of the numbered cell. Traces are vertically offset for clarity. Scale bars: 5 ms, 20 mV. Cell 1: □L = 40.0 ± 3.1%, p < 0.01. Cell 2: □L = 0.0 ± 3.5%, p = 0.70. Cell 3: □L = -0.1 ± 3.4%, p = 0.34. Cell 4: □L = 29.1 ± 3.0%, p < 0.01 (Wilcoxen signed-rank tests). **D - F:** Representative heterocellular network among SOM^-^ and SOM^+^ neurons, as described for **A-C.** Area 0: □L =7.0 ± 3.1%, p = 0.11. Cell 1: □L = 22.6 ± 1.8%, p < 0.01. Cell 2: □L = 0.0 ± 2.1%, p = 0.56. Cell 3: □L = 7.3 ± 1.0%, p = 0.03 (Wilcoxen signed-rank tests). **G – I:** Example homocellular network among PV^+^ neurons, as described for **A-C**. Cell 1: □L = 45.8 ± 0.9%, p < 0.01. Cell 2: □L = 13.8 ± 2.4%, p < 0.01. Cell 3: □L = 52.0 ± 0.9%, p < 0.01. Cell 4: □L = 4.2 ± 1.9%, p = 0.08 (Wilcoxen signed-rank tests).

**Fig. 4.**
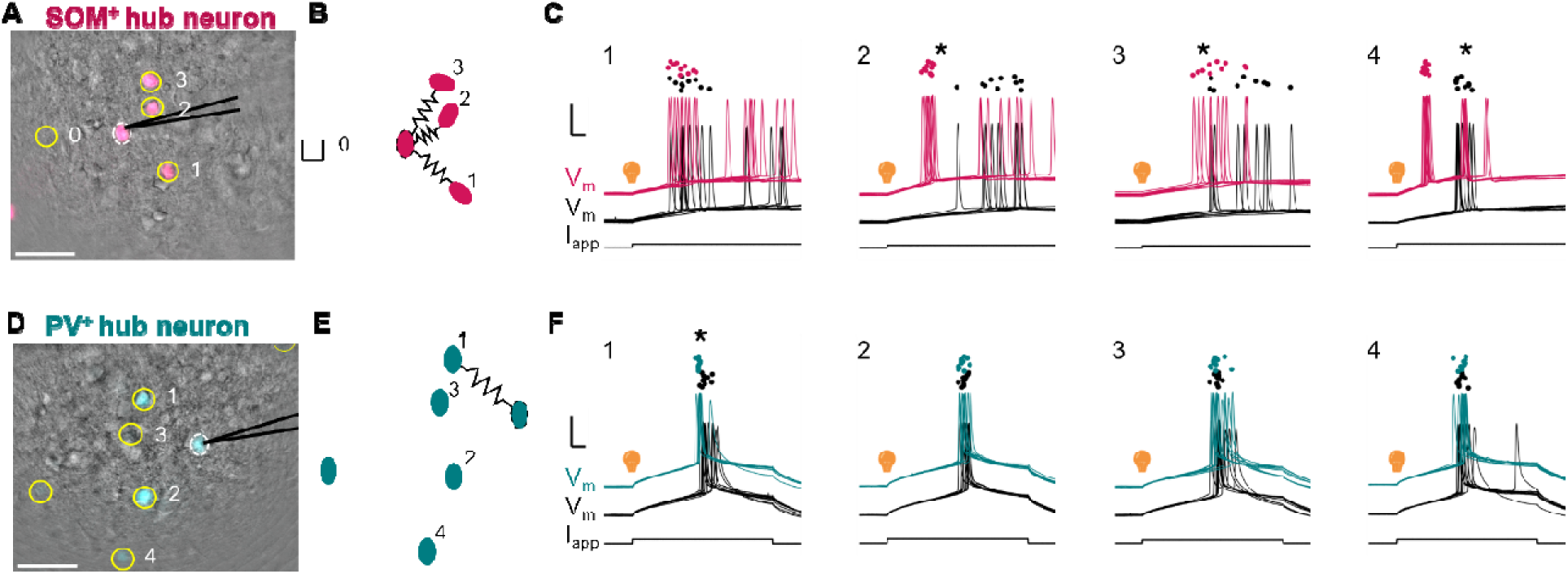
Maps of coupled networks in cortex. **A.** Overlay of IR and fluorescence images from live recording of st-ChroME-mRuby-expressing SOM^+^ somatosensory cortical neurons *in vitro*; fluorescence is pseudocolored magenta. The patched neuron is circled with white dashes. Cells tested by opto-□L are circled in yellow. One empty control area is marked by a square. Scale bar: 50 μm. **B.** Diagram of the coupled network embedded within all tested neurons from **A**. **C.** Spiking responses from opto-□L measurements from numbered cells in **A**. Dots mark the times of first spikes in the patched neuron from each trial. Black traces are responses of the patched neuron during rheobase-only trials and colored traces are responses during added focal photostimulation of the numbered cell. Traces are vertically offset for clarity. Scale bars: 5 ms, 20 mV. Area 0: □L = 10 ± 5.4%, p = 0.13 (Wilcoxon signed-rank test). Cell 1: □L = 63.0 ± 1.1%, p < 0.01. Cell 2: □L = 35.3 ± 5.0%, p = 0.01. Cell 3: □L = 55.8 ± 1.1%, p < 0.01. **D - F:** Coupled network among cortical PV^+^ neurons., as described for **A-C.** Cell 1: □L = 9.0 ± 0.6 %, p < 0.01, Cell 2: □L = 0.0 ± 1.5%, p = 0.25 Cell 3: □L = 0.0 ± 2.7%, p = 0.54, Cell 4: □L = 0.0 ± 2.0%, p = 0.49. Six additional more-distant neurons without electrical synapses were tested for this hub neuron that are not shown.

To extend opto-δL as a tool for measuring and mapping coupled networks across the brain, we used opto-δL to map coupled networks in primary somatosensory cortex. In 1 slice and 1 patched hub cell from a mouse expressing st-ChroME in SOM^+^ neurons (Fig. 4A-C), we identified one cortical network that comprised three neurons with δLs of 63.0 ± 1.1%, 35.3 ± 5.0%, and 55.8 ± 1.1%. The average δL in the network was 51.4 ± 8.3% and average intersomatic distance was 43.8 μm. In 1 slice and 1 patched hub cell from a PV-Cre mouse (Fig. 4D-F), we identified one cortical network that was limited to two neurons, with a δL of 9.0 ± 0.6 and intersomatic distance of 43.5 μm. General conclusions regarding cortical coupled networks are limited from these two examples, but they demonstrate the potential broad applicability of the method. Opto-δL can also be measured using 2-photon photostimulation (Supp. Fig 5).

**Fig. 5.**
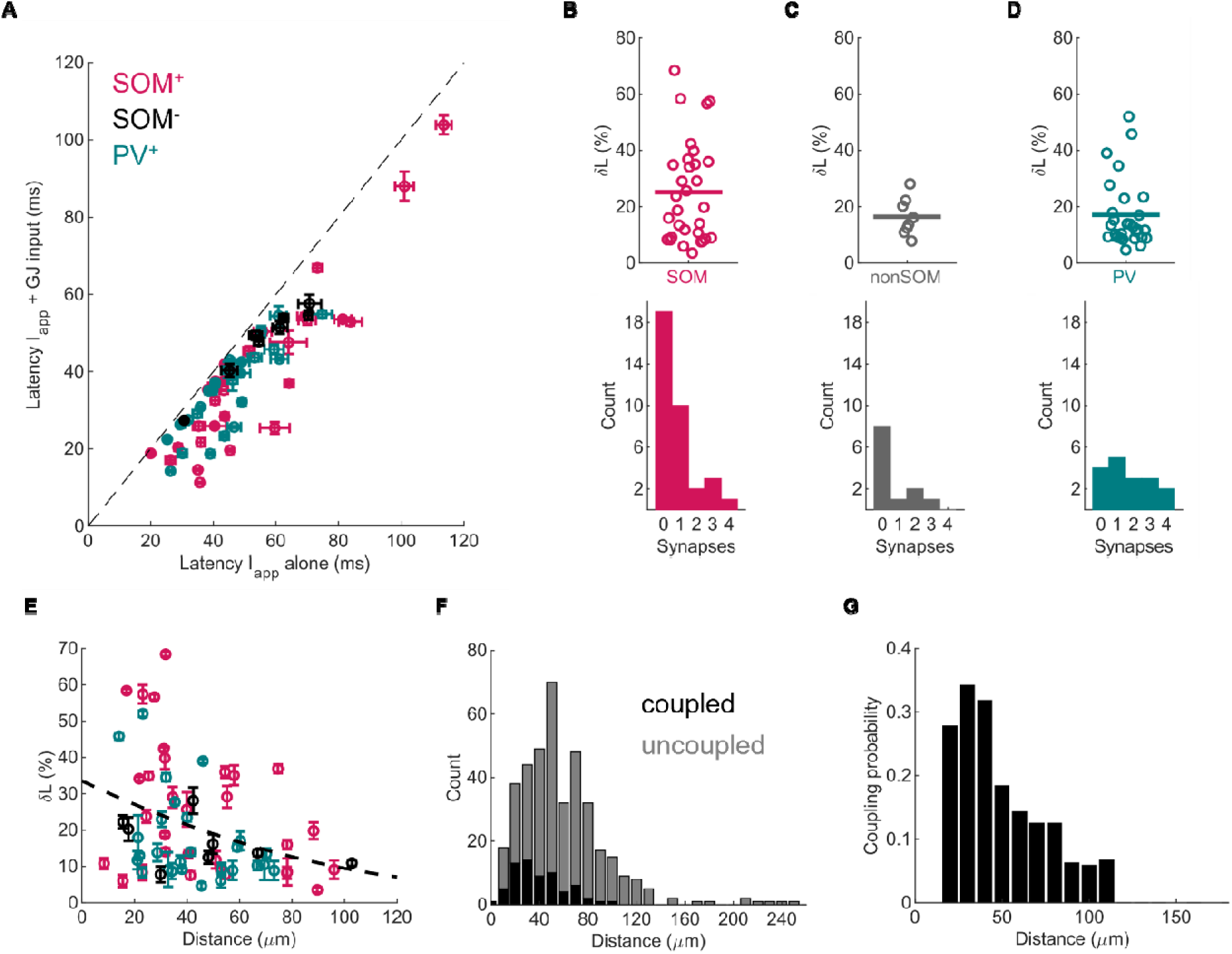
Distribution of TRN electrical synapses and networks over strength and space. **A.** Latencies (± SEM) used to compute □L emanating from 16 SOM^+^, 4 SOM^-^ and 13 PV^+^ hub neurons. **B.** Top: □L for homocellular SOM^+^ synapses (mean □L = 25.0 ± 3.2%, n = 31 synapses from 16 networks in 19 slices from 19 mice. Bottom: distribution of network sizes (number of coupled neighbors per hub cell) for SOM^+^/SOM^+^ networks (mean size: 1.7 ± 0.1 synapses per map). **C.** As in **B,** for heterocellular SOM^-^/SOM^+^ synapses (mean □L = 16.4 ± 2.4%, n = 8 synapses from 4 networks in 4 slices from 4 mice; mean size 2.0 ± 0.2 synapses per network). **D.** As in **B** for homocellular PV^+^ synapses (mean □L = 17.1 ± 2.3%, n = 28 synapses from 13 networks in 13 slices from 12 mice; mean size 2.2 ± 0.1 synapses per network, n = 13 maps). **E.** Distribution of □L across intersomatic distance for each subtype in TRN (n = 67 synapses). **F.** Number of photostimulated neurons that were (black, n = 67) or were not (grey, n = 331) coupled to a patched hub neuron, plotted against distance from the hub neuron (n = 398 total connections tested). **G.** Probability of coupling versus intersomatic distance.

To fully characterize mature TRN networks, we used opto-□L to test connections emanating from 70 patched hub neurons (mean R_in_ 150.4 ± 7.8 MΩ, mean resting V_m_ -68 mV ± 0.7 mV) from 16 slices taken from 15 PV-Cre mice and 49 slices taken from 46 SOM-Cre mice injected with AAV9-CAG-DIO-ChroME-st-P2A-H2B-mRuby into the TRN, ultimately testing 398 possible connections between hub neurons and neighboring neurons that revealed 33 networks (16 SOM^+^, 4 SOM^+^/ SOM^-^, 13 PV^+^). For each hub neuron, we tested an average of 6.3 neighboring neurons for electrical synapses; the largest number tested was 20 neighboring neurons. Overall, we measured 67 electrical synapses via opto-□L across all combinations of genetic subtypes tested (Fig. 4A). Sampling 40 SOM^+^ hub neurons (mean R_in_ 171.2 ± 11.2 MΩ, mean resting V_m_ -68 mV ± 1.0 mV), we found 31 synapses between SOM^+^ hub neurons and SOM^+^ neighbors, ultimately mapping 16 homocellular networks (36 slices from 33 mice). The mean coupling incidence was 15.1% across all tested SOM^+^ connections (n = 206 connections tested). Some TRN cells appear to lack Cx36 expression (*58*); limiting our computation of coupling incidence to hub cells that had at least one synapse, we found coupling incidence was 40.3% (n = 67 possible connections). The mean □L between SOM^+^ neurons was 25.0 ± 3.2%, and the mean homocellular SOM network contained 1.7 ± 0.1 electrical synapses per hub neuron (Fig. 4B). For heterocellular networks, we patched 13 patched SOM^-^ hub neurons (mean R_in_ 115.6 ± 14.3 MΩ, mean resting V_m_ -69 mV ± 2.1 mV) and photostimulated neighboring SOM^+^ neurons (Fig. 3C); we measured 8 significant values of □L, ultimately mapping 4 heterocellular networks (13 slices from 12 mice). The mean coupling rate was 7.3% over all 85 connections tested and 25.8% across 31 tested connections from a hub cell with at least one connection. The mean □L for SOM^+^ to SOM^-^ connections was 16.4 ± 2.4%, and the mean heterocellular network contained 2.0 ± 0.2 direct electrical synapses per hub neuron (Fig. 4C). For homocellular networks between PV^+^ and PV^+^ neurons, we recorded from 17 PV^+^ hub neurons (mean R_in_ 128.3 ± 10.3 MΩ, mean resting V_m_ -68 mV ± 1.3 mV) and measured 28 synapses using opto-□L, ultimately mapping 13 homocellular networks (16 slices from 15 mice). The mean coupling rate was 25.9% across all 108 tested connections, and 39.1% from hub cells with least one electrical synapse (n = 74 connections tested). The mean □L for PV^+^ to PV^+^ electrical synapses was 17.1 ± 2.3% and the mean network size was 2.2 ± 0.1 direct electrical synapses per hub neuron (Fig. 4D). Across networks, there were no significant differences in □L between subtypes (Fig. 4B-4D, p > 0.05). The mean □L across all subtypes was 22.8% ± 1.9% (n = 67 synapses), and 50% of synapses had □L between 11.0% and 33.3%. The largest □L was 68.4%. It was rarer to fail to find electrical synapses between PV^+^ neurons (Fig. 4B-D, p < 0.01, chi-squared test). PV^+^ hub neurons had no electrical synapses to nearby opsin-expressing cells in 23.5% of recorded neurons, while SOM^+^ hub neurons had no connections 54.3% of the time, and SOM^-^ hub neurons had no connections 66.7% of the time. There was no difference between SOM^+^ and SOM^-^ hub neurons in the frequency of cells that had no nearby electrical synapses (p = 0.07, chi-squared test).

We also used opto-δL to more exhaustively and precisely sample the spatial extent of coupling from each hub neuron. Measuring electrical synapses by patching pairs introduces spatial selection biases to the pairs that are ultimately tested in experiments; this effect may have contributed to a discrepancy between results from rat TRN suggesting that single electrical synapses occur only within intersomatic distances less than 40 μm (*75*), and results from dye-coupling, which permits spread of dye over several synapses, indicating a median intersomatic distance of 80-120 μm (*64*). We leveraged the ability of opto-□L to stimulate distant neurons, testing synapses up to 250 µm. Across all neuron types grouped together, we report that TRN neurons that were closer together were coupled more strongly (Fig. 4E) and more likely to be coupled (Fig. 4F-G), confirming previous reports that coupling diminishes with intersomatic distance. Collectively, our data also showed that electrically coupled networks for all cell types link neurons as far as 100 μm apart (Fig. 5E-G). Taken together, our measurements offer new insight into the precise form and functions of the coupled networks of the mature mammalian TRN.

## Discussion

Connexin36 is widely expressed throughout the mature mammalian brain, yet functional measurements and maps of electrically coupled networks remain absent in live tissue. Here, we introduce a novel method to map networks in living tissue, opto-□L, which leverages focal photostimulation of soma-targeted opsin-expressing neurons combined with measurements of changes in spike timing, including tests of significance, that result from excitation across electrical synapses. Using opto-□L, we show that GABAergic TRN neurons of differing molecular subtypes are coupled in homocellular and heterocellular electrical networks and extend further across tissue than previously identified by dual-cell patching (*75*). Further, we identify functional network size, which is limited to far fewer neurons than dye coupling had previously suggested (*64*). These results imply that the electrical synapses and networks of the mature TRN provide a link between thalamocortical pathways, and that they do so by forming networks that are selective in size.

In our experiments in TRN, we observed PV^+^ neurons without electrical synapses less frequently than among SOM^+^ neurons (Fig. 5B). We hypothesize that is because PV expression is more of an umbrella than a distinct marker for a subset of TRN neurons, as has long been believed (*54, 55, 80, 82–85*), and therefore, our maps of PV^+^ neurons might include both heterocellular and homocellular electrical synapses. However, other experiments suggest that PV^+^ neurons are a distinct subset of TRN neurons that primarily form connections with first-order thalamus (*53, 86*), similar to subsets identified by the protein CB (*55*) and the transcription factor SPP1 (*54*), and that they are distinct from SOM^+^ neurons. If the latter were the case, our results imply that TRN neurons that project to first-order thalamus form more electrical synapses than SOM^+^ neurons. In turn, that difference would imply that coordination between larger numbers of TRN neurons is more prevalent for first-order thalamus.

The electrical connections we observed as networks emanating from one neuron were small, ranging from 1 to 4 electrical synapses; these are the first direct measurements of the number of neurons composing a living coupled network. A previous dye-coupling study in juvenile TRN stained networks with an average of 7 neurons, and as large as 10 to 20 neurons (*64*). Dye-coupling allows spread of dye between neurons that are not directly coupled to the originally patched neuron, and thereby less likely to influence spike its times. In contrast, opto-δL is likely to detect only direct connections, as it is quite unlikely that an indirectly coupled neuron would affect spike timing by the magnitudes that we regularly observed. The overall proportion of neurons that were coupled in our measurements was 16.8%, which is less than that reported from paired recordings in juvenile rats and mice, where reports vary between 31% (*58*), 45% (*56*), and 71% (*75*). There are several possible explanations for this discrepancy. The most likely cause for this difference is that our tests and maps included TRN neurons that were farther away from the patched neuron, while previous paired-patching experiments often select for neurons that are much closer together. Species differences or decreases in coupling frequency from juveniles to adults could also contribute to the discrepancy. Our observations of networks composed of 1-4 local neurons are consistent with results inferring that TRN neurons form small clusters of coupled neurons (*75*). We interpret the small sizes of these networks to indicate that electrical coupling and network membership are more selective and targeted than ubiquitous or promiscuous. The full extent of networks beyond a single hub neuron, the specific rules that determine connectivity, and the functional outcomes for thalamocortical processing, remain open and important questions.

Our experiments likely undersampled electrical synapses and networks for several reasons. First, slicing brain tissue cuts some connections, causing some neuron loss. Second, our previous experiments determined that heterocellular synapses are common (*56*), but we were unable to photostimulate neurons that did not express opsin, which represent another pool of possibly undetected synapses. Failure of virus transfection for some neurons could be a source of misidentification in our experiments. Furthermore, due to finite recording times, we only tested a subset of nearby neurons for each hub neuron; it is possible that we missed some untested connections in experiments. Our method may also underestimate connections due to a floor effect from our light scatter compensation and differences in sensitivity between cc and □L to detect very weak synapses. Therefore, while our experiments failed to detect any electrical synapses for 31 neurons, we expect that we overestimated the number of TRN neurons without electrical synapses. It is possible that a subset of TRN neurons exists without electrical synapses, which would be consistent with the observation that some TRN neurons do not stain for Cx36 (*58*). Likewise, we interpret our report of 1 to 4 electrical synapses for TRN networks as establishing a floor for network size.

Opto-□L can be implemented in other brain regions, as demonstrated by the network maps we created in cortex (Fig. 3). Unlike TRN, cortex is a mix of inhibitory and excitatory neurons, and coupled inhibitory neurons may be separated by longer distances. In cortex, an additional barrier to identifying and measuring coupling by paired patching is the need to visually identify GABAergic neurons. Use of fluorescent labels in opto-□L mitigates this barrier for homocellular networks. Measurements of □L can in principle be used *in vivo* with focal holographic stimulation (*74, 87*) to map electrical synapses, and we suggest that this would be a valuable next step to more accurately map TRN networks fully in three dimensions. Opto-δL opens possibilities for more easily detecting and measuring coupled networks throughout the brain. For example, one could examine how network membership and dynamics change in response to different plasticity paradigms. Combining opto-δL with activity indicators may reveal how activity propagates through a network under varied conditions.

Opto-□L and cc are correlated (Fig. 1D), but due to the nonlinearities of spike generation, there is no simple translation between the two quantities. Both methods have limitations. Measurement of cc is modulated by subthreshold nonlinearities, which can be amplified by as much as a factor of 4 by the persistent sodium current (*12, 88, 89*) and is similarly modulated by other voltage-dependent conductances that are active near rest (*90–93*) (reviewed in Pereda et al. 2013 (*94*)). By the same tokens, cc is inexorably intertwined with resting membrane potential, the amplitude of injected current used to measure it, and changes with drift or any other factor that alters input resistance. Electrical synapses often express full or partial rectification; cc detects rectification readily, while opto-□L would require separately patching both neurons of an identified pair and taking care to match initial latencies, to measure asymmetries using opto-□L. In addition to light scatter (discussed in Methods), opto-□L is also, for the reasons above, sensitive to resting membrane potential and the magnitude of spike-driving currents used (*73*). Smaller amounts of current that yield correspondingly longer initial latencies allow for more robust measurements, but spiking precision diminishes as one approaches spiking threshold. Therefore, current application above rheobase, but only enough so to drive reliable spikes, is ideal for □L measurements. In comparing cc and opto-□L, it must be noted that opto-□L clarifies the functional impact of an electrical synapse – that is, electrical synapses shift spike times, often by substantial amounts, in the neurons and networks they couple.

## Materials and Methods

### Animals

Experiments were approved by the Lehigh University IACUC. SOM-Cre (Jax: 013044) or PV-Cre (Jax: 008069) mice were used for experiments. Mice were at least P28 for surgeries and P42 for electrophysiological recordings. For one juvenile experiment (Fig. 1), a calbindin-Cre (Jax: 028532) mouse was injected at P0 and electrophysiological recordings occurred at P14. To target our CRISPR knock down selectively to the PV-expressing GABAergic neurons we crossed male Rosa26-lox-stop-lox-Cas9/GFP mice (Jax: 026175) with PV-Cre (Jax: 017320), generating Cas9 and GFP reporter expression in PV neurons (PV-Cre-CRISPRCas9-GFP) (*95*).

### Electrophysiology

Mice were transcardially perfused with NMDG ACSF solution (in mM): 92 NMDG, 2.5 KCl, 1.25 NaH_2_PO_4_, 30 NAHCO_3_, 20 HEPES, 25 glucose, 2 thiourea, 5 NA ascorbate, 3 NA pyruvate, 0.5 CaCl_2_, and 10 MgSO_4_·7H_2_O (315 mOsm L^−1^, 7.3-7.4 pH). Horizontal brain slices 250 µm thick were cut and incubated in NMDG solution. Slices were incubated at 34°C for 12-15 min in NMDG following cutting and returned to HEPES ACSF at room temperature until recording. The HEPES ACSF solution comprised in mM: 92 NaCl, 2.5 KCl, 1.25 NaH_2_PO_4_, 30 NaHCO_3_, 20 HEPES, 25 glucose, 2 Thiourea, Na ascorbate, NA pyruvate, 2 CaCl2, and 2 MgSO_4_·7H_2_0 (315 mOsm L^−1^, 7.3-7.4 pH). The ACSF bath during recording contained (in mM): 126 NaCl, 3 KCl, 1.25 NaH_2_PO_4_, 2 MgSO_4_, 26 NaHCO_3_, 10 dextrose, and 2 CaCl_2_ (315–320 mOsm L^−1^, saturated with 95% O_2_/5% CO_2_). The submersion recording chamber was held at 34°C (TC-324B, Warner Instruments). Electrodes were filled with (in mM): 135 potassium gluconate, 2 KCl, 4 NaCl, 10 Hepes, 0.2 EGTA, 4 ATP-Mg, 0.3 GTP-Tris, and 10 phosphocreatine-Tris (pH 7.25, 295 mOsm L^−1^). 1 M KOH was used to adjust pH of the internal solution. The approximate bath flowrate was 2 ml min^−1^ and the recording chamber held approximately 5 ml solution.

The TRN was visualized under 4x magnification, and pairs of adjacent TRN cells from any sensory sector were identified and patched under 40x IR-DIC optics (SliceScope, Scientifica, Uckfield, UK). Cell type was determined by mRuby epifluorescence in the patched neurons. Cells with low fluorescent contrast were avoided. The mRuby reporter was excited by a 560 nm diode delivered through the objective (CoolLED pE-300). Voltage signals were amplified and lowpass filtered at 8 kHz (MultiClamp, Axon Instruments, Molecular Devices, Sunnyvale, CA, USA), digitized at 20 kHz with custom Matlab routines controlling a National Instruments (Austin, TX, USA, USB6221 DAQ board), and data were stored for offline analysis in Matlab (Mathworks, R2023b, Natick, MA, USA). Recordings were made in whole-cell current-clamp mode. Negative current was used to maintain cells at −70 mV to -60 mV during recordings. Pipette resistances were 5 - 9 MΩ before bridge balance; recordings were discarded if access resistance exceeded 25 MΩ. Voltages are reported uncorrected for the liquid junction potential.

### Stereotaxic surgery

SOM-Cre (Jax: 013044), PV-Cre (Jax: 008069), or PV-Cre-CRISPR-Cas9-GFP mice were anesthetized via inhaled isoflurane (0.8-1.2%) and 0.1 mL of 0.015 mg/mL buprenorphine via intraparietal injection. Mice were positioned on a heated pad and in a stereotax (Kopf). After exposure of the skull, a dental drill was used to open a hole above the injection site. Then, AAV9-CAG-DIO-ChroME-st-P2A-H2B-mRuby (titer 2.7 x 10^13^ GC/mL) was injected at a rate of 100 nL/min (WPI UMP-3). Viral vectors used for knocking down the Cx36 or Cx29 were designed by David Uygun and Radhika Basheer as described previously (*95*). Briefly, the PX552 plasmid (Addgene plasmid #60958) was modified to encode triple U6-sgRNA cassettes each targeting a distinct locus of the target genes. The GFP sequence was replaced by the sequence encoding the blue fluorescent protein (BFP). The coordinates for injections were -1.35 AP, ± 2.1 ML, -3.0 and -3.2 DV for TRN and -0.5 AP, ± 3.0 ML, -2.0 and -2.6 DV for somatosensory cortex. We injected a total of 600 nL of virus for TRN and 800 nL of virus for somatosensory cortex. Post surgery and scalp suturing, mice were given an intraperitoneal injection of 0.1 mL of 0.5 mg/mL carprofen and 1 mL saline, then monitored for recovery. Mice were given at least two weeks for viral expression before experiments. For the juvenile surgery, a calbindin-Cre (Jax: 028532) mouse was anesthetized via hypothermia on a cold metal plate, and the coordinates for injection targeting TRN were -1.4 AP, ± 1.9 ML, -1.4 and -1.9 DV.

### Image processing

All fluorescence images were modified in Fiji and pseudocolored to facilitate visualization. All infra-red differential interference contrast images were flat-field corrected.

### Opto-□L

For the rheobase current necessary to measure □L, we used the smallest stimuli that drove spiking at least 80% of the time, with the CV of latency less than 0.2. Trials were excluded if latency was less than 20 ms when the neuron was given a rheobase stimulus.

To perform single-cell optogenetic stimulation, we used a 560 nm LED (Mightex) and digital mirror device (Mightex Polygon 1000) to direct light to an area the size of an individual soma (7 - 8 µm diameter circle). The area used covered the entire soma without extending beyond the edge of the soma. Prior to any experiments, we patched an opsin-expressing cell to determine an intensity of light that was sufficient to drive bursts for a 50 ms stimulus, and we then used a photostimulus that was 50% stronger for all subsequent experiments in that slice. Light intensity ranged from 3.7 - 99.8 mW/mm^2^ with most experiments occurring in 6.2 - 25.0 mW/mm^2^ range.

Single-cell optogenetic stimulation is confounded by off-target excitation by light scatter for both 1 photon and 2 photon stimulation paradigms (*76–78*). For the purposes of mapping synaptic connections, scatter-induced photostimulation of the patched neuron is a potential confound. Depolarization from photostimulation caused by light scatter is directly proportional to the distance between the patched neuron and focal stimulation (Supp. Fig. 3). Opto-δL is well-suited to overcome the problem of unintended photostimulation to the patched neuron because the method quantifies latency of spikes rather than voltage changes. Consequently, comparing latency when a nearby neuron is photostimulated to latency when an equidistant empty control location is photostimulated allowed us to distinguish the contribution of GJs to spike times.

After patching TRN neurons from any sensory sector, we used one of two methods to first make measurements that would control for the effects of light scattering and then to compute δL.

i) Direct corrections to δL: We delivered a rheobase stimulus to the patched neuron alone 10 times. Interleaved between those stimulations, we gave the neuron a rheobase stimulus while also focally photostimulating either a nearby mRuby^+^ neuron or an equidistant empty spot. Offline, we measured the latency of spikes from stimulation onset for each condition. We report □L as the change in latency induced by focal photostimulation of the nearby neuron minus the change in latency induced by photostimulation of an empty area.
ii) Correction to Iapp: We applied focal photostimulation ten times to an empty location equidistant from the patched neuron as the nearby neuron of interest. We measured the average depolarizing current that resulted from the light scattering. We then adjusted rheobase Iapp by average current caused by light scatter (Supp. Fig. 4) and used that adjusted Iapp when photostimulating the nearby mRuby^+^ or an equidistant empty location. Offline, we calculated □L by comparing latency changes between when light targeted the empty control location 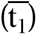 compared to light targeting the mRuby^+^ neuron 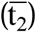.

We used directly-corrected δL for 27 homocellular network measurements, and we used correction to I_app_ for 30 homocellular network measurements.

### Numerical analysis and statistics

To calculate δL we used the average spike latency from control trials (*t_1_*) and average spike latency when a nearby neuron was photostimulated (*t_2_*):

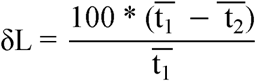

We tested δLs for significance with a Wilcoxon signed-rank test. We compared δLs between different neuronal subtypes via Mann-Whitney U test, and we used Chi-squared tests to compare electrical synapses counts per network between different genetic subtypes. Statistical tests were considered significant when p < 0.05. In our analysis of network size, we only included experiments where three or more possible connections were tested.

## Acknowledgments

We thank all members of the Haas and Bender laboratories and Alberto Pereda for fruitful discussions.

## Funding

NIH NS128713 and Whitehall Grant in Aid (JSH),

Veterans Administration Merit Award I01BX006105 (RB)

NINDS R01 NS119227 (RB)

VA CDA2 award IK2 BX004905 (DSU)

## Author contributions

Conceptualization: JSH

Methodology: MJV, JSH, KJB, DSU, RB

Investigation: MJB, JSH

Writing—original draft: MJV, JSH

Writing—review & editing: MJV, DSU, RB, KJB, JSH

## Data and materials availability

Data are available upon request from the corresponding author.

## Table of contents for supplementary material

**Fig. S1. Validation of single-cell focal photostimulation.**

**Fig. S2. Additional maps of coupled networks between TRN neurons**

**Fig. S3. Compensation for effects of scatter-induced photoexcitation on spike times.**

**Fig. S4. Example of excitation from light scattering.**

**Fig. S5. Opto-**δ**L mapping of a coupled network with 2-photon stimulation.**

