## Supplemental_Data for "Photomapping electrically coupled networks of the mature thalamus and cortex"

Science Advances

AAAS

Mitchell Vaughn *et al.* For papers with only two authors:

**This PDF file includes:**

Figs. S1 to S5

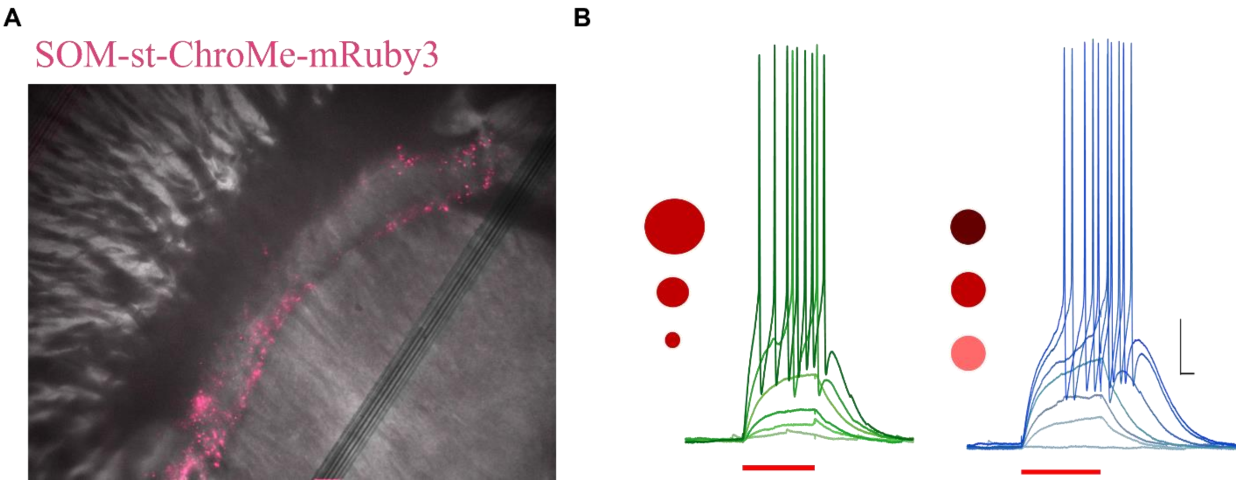


**Fig. S1. Validation of single-cell focal photostimulation.**

**A.** Expression pattern of st-ChoME-mRuby3 via AAV injection in a SOM-Cre mouse (fluorescence pseudocolored magenta). Expression was restricted to the shell of TRN.
**B.** Left, spiking driven by increasing area of light in a spot (3.14, 7.07, 14.14, 22.0, 31.4, 47.2 µm2 circles). Right, spiking driven by increasing light intensity (24.4, 37.1, 59.2, 66.2, 80.1, 87.0, 105.7 mW/mm2). Scale bar 25 ms, 10 mV. Vm = -61 mV.

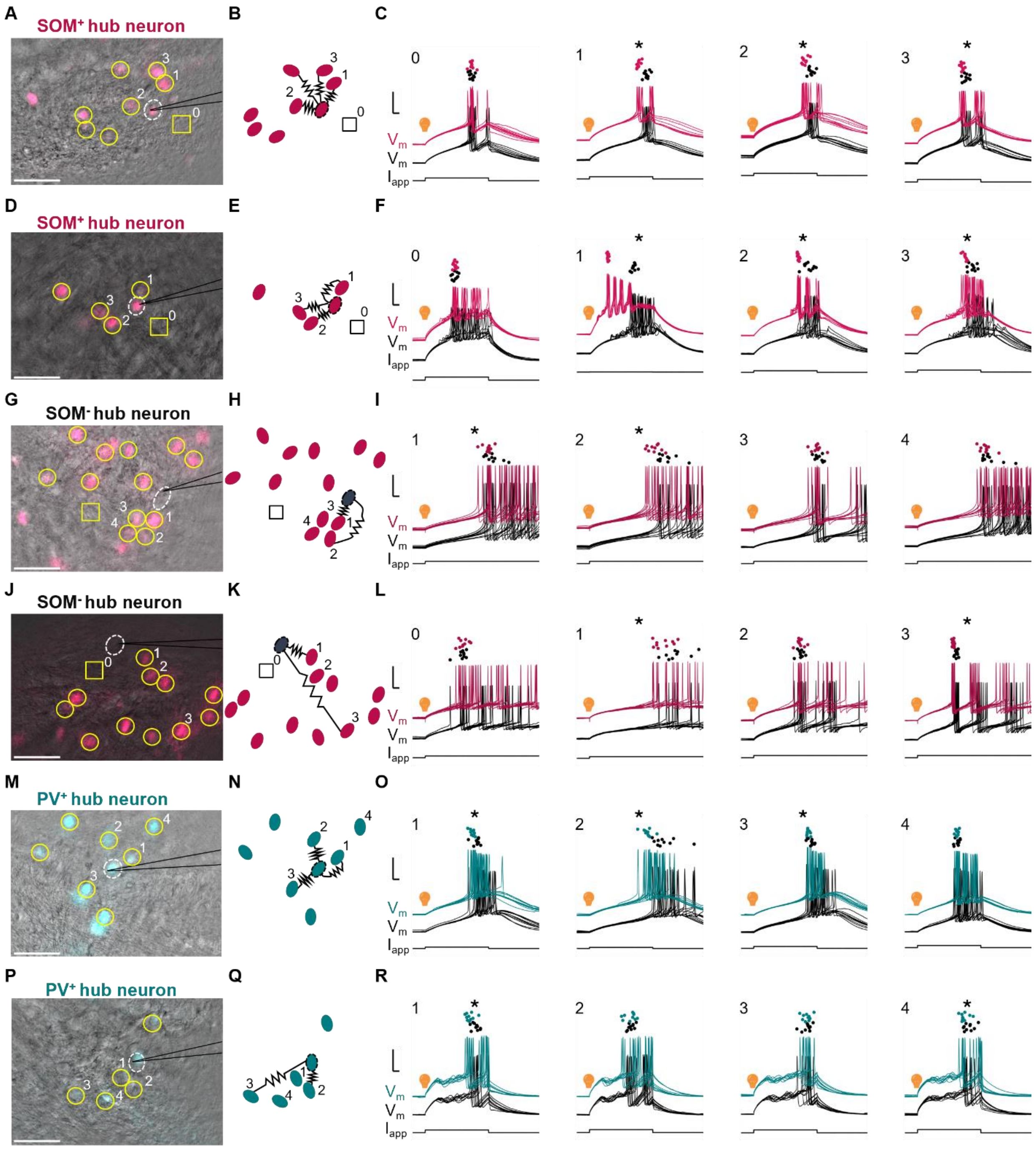


**Fig. S2. Additional maps of coupled networks between TRN neurons**

**A.** Overlay of IR and fluorescence images from live recording of st-ChroME-mRuby-expressing $SOM^+$ TRN neurons *in vitro*; fluorescence is pseudocolored magenta. The patched neuron is circled with white dashes. Cells tested by opto-δL are circled in yellow. One empty control area is marked by a square. Scale bar: 50 μm.

**B.** Diagram of the coupled network embedded within all tested neurons from **A**.

**C.** Spiking responses from opto-δL measurements from numbered cells in **A**. Dots mark the times of first spikes in the patched neuron from each trial. Black traces are responses of the patched neuron during rheobase-only trials and colored traces are responses during added focal photostimulation of the numbered cell. Traces are vertically offset for clarity. Scale bars: 5 ms, 20 mV. Area 0**:** $\delta L = 0.0 \pm 1.6\%$, $p = 0.98$. Cell 1: $\delta L = 14.0 \pm 1.0\%$, $p < 0.01$. Cell 2: $\delta L = 10.8 \pm 1.5\%$, $p < 0.01$. Cell 3: $\delta L = 7.5 \pm 1.1\%$, $p < 0.01$ (Wilcoxen signed-rank test).

**D - F:** Representative homocellular network among $SOM^+$ neurons, as described for **A-C.** Area 0: $\delta L = 0.3 \pm 1.8\%$, $p = 0.92$. Cell 1: $\delta L = 58.4 \pm 0.5\%$, $p < 0.01$. Cell 2: $\delta L =18.7 \pm 0.5\%$, $p < 0.01$. Cell 3: $\delta L = 13.5 \pm 1.1\%$, $p < 0.01$ (Wilcoxen signed-rank test).

**G - I:** Representative heterocellular network among $SOM^-$ and $SOM^+$ neurons, as described for **A-C.** Cell 1: $\delta L = 7.8 \pm 2.2\%$, $p = 0.02$. Cell 2: $\delta L = 16.1 \pm 2.6\%$, $p < 0.01$. Cell 3: $\delta L = 3.3 \pm 2.5\%$, $p = 0.45$. Cell 4: $\delta L = 7.5 \pm 2.6\%$, $p = 0.11$ (Wilcoxen signed-rank test).

**J - L:** Representative heterocellular network among $SOM^-$ and $SOM^+$ neurons, as described for **A-C.** δL: Area 0: $\delta L = 0.0 \pm 4.0\%$, $p = 0.5$. Cell 1: $\delta L = 17.4 \pm 3.2\%$, $p = 0.03$. Cell 2: $\delta L = 0.0 \pm 2.6\%$, $p = 0.70$. Cell 3: $\delta L = 8.1 \pm 0.8\%$, $p < 0.01$ (Wilcoxen signed-rank test).

**M – O:** Example homocellular network among PV+ neurons, as described for **A-C**. Cell 1: $\delta L = 8.4 \pm 1.4\%$, $p = 0.03$. Cell 2: $\delta L = 22.9 \pm 2.1\%$, $p < 0.01$ Cell 3: $\delta L = 4.6 \pm 0.8\%$. $p = 0.04$, Cell 4: $\delta L = 0.0 \pm 1.2\%$, $p = 0.63$ (Wilcoxen signed-rank test).

**P – R:** Example homocellular network among PV+ neurons, as described for **A-C**. δL: Cell 1: $\delta L = 11.7 \pm 2.5\%$, $p < 0.01$. Cell 2: $\delta L = 0.0 \pm 4.2\%$, $p = 0.77$. Cell 3: $\delta L = 0.0 \pm 2.5\%$, $p = 0.39$, Cell 4: $\delta L = 8.8 \pm 2.6\%$, $p = 0.04$ (Wilcoxen signed-rank test).

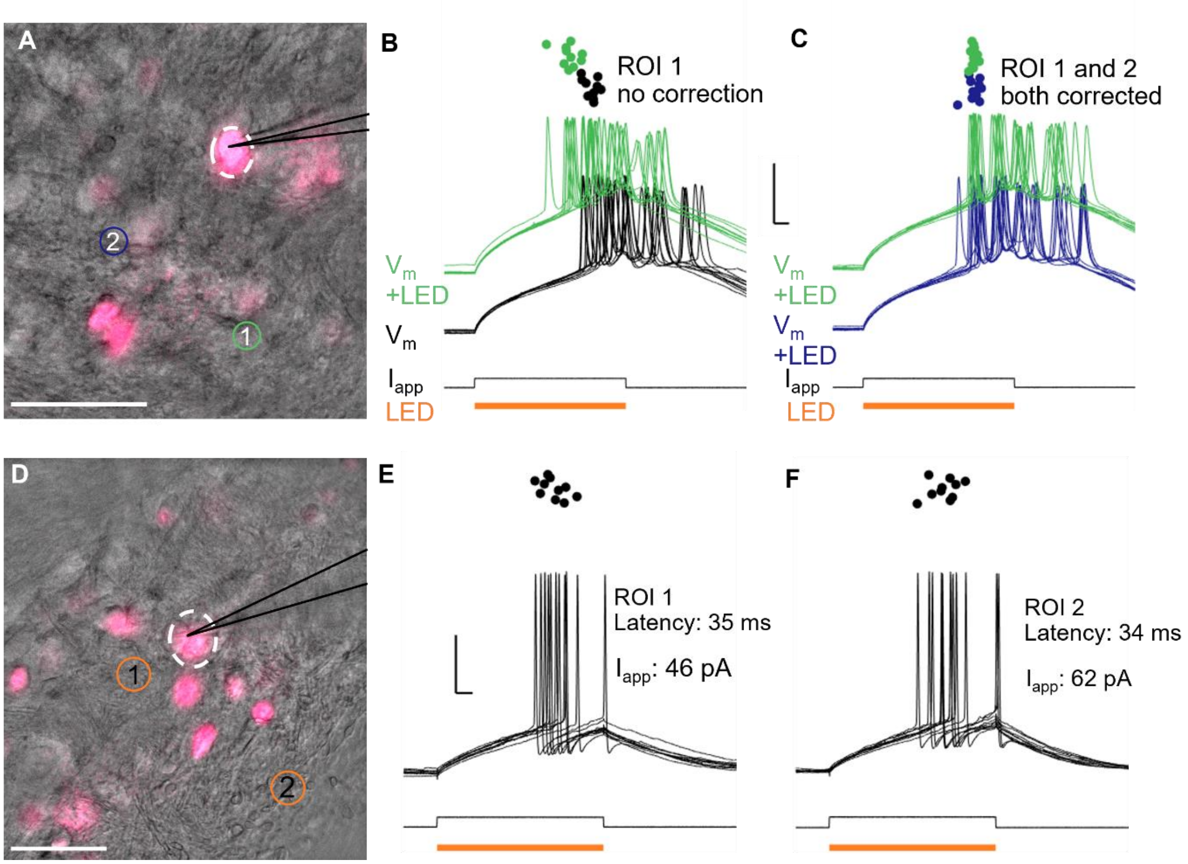


**Fig. S3. Compensation for effects of scatter-induced photoexcitation on spike times.**

**A.** Overlay of IR and fluorescent images of ChroME-mRuby-expressing neurons in a SOM-Cre mouse. Fluorescence is pseudocolored magenta. Scale bar: 50 µm. The patched neuron is circled with white dashes. Empty regions of interests (ROIs) (e.g. orange circles) at different distances were photostimulated to determine the compensation for scatter to rheobase for the patched neuron.

**B.** Spiking responses for rheobase (51 pA) applied current alone (black traces; mean latency 38.6 ms) or during simultaneous photostimulation of ROI1 (green traces, mean latency 31.2 ms). Dots mark the times of first spikes from each trial. Uncompensated scatter from ROI 1 caused a false δL of 19%.

**C.** Spiking responses for compensated rheobase (22 pA) and simultaneous photostimulation of ROI 2 (blue traces; mean latency 36.6 ms), and for compensated rheobase during photostimulation of ROI 1 (green traces; mean latency 36.5 ms).

**D**. Overlay of IR and fluorescence (pseudocolor: magenta) images of ChroME-mRuby-expressing neurons in a slice from a SOM-Cre mouse. Scale bar 50 µm. The patched neuron is circled with white dashes. Without any photostimuli, rheobase was determined to be 65 pA. Two non-fluorescing ROIs, circled in orange, were selected for photostimulation.

**E.** Spiking in the recorded neuron after correction of rheobase (square pulse, lower) for scatter to 46 pA during photostimulation of ROI 1 (orange bar). The corrected mean latency of the first spike was 35 ms.

**F.** Spiking in the recorded neuron after correction of rheobase for scatter to 62 pA (square pulse, below) during photostimulation of ROI 2 (red bar). The mean corrected latency of the first spike was 34 ms. Scale bars are 25 ms, 10 mV.

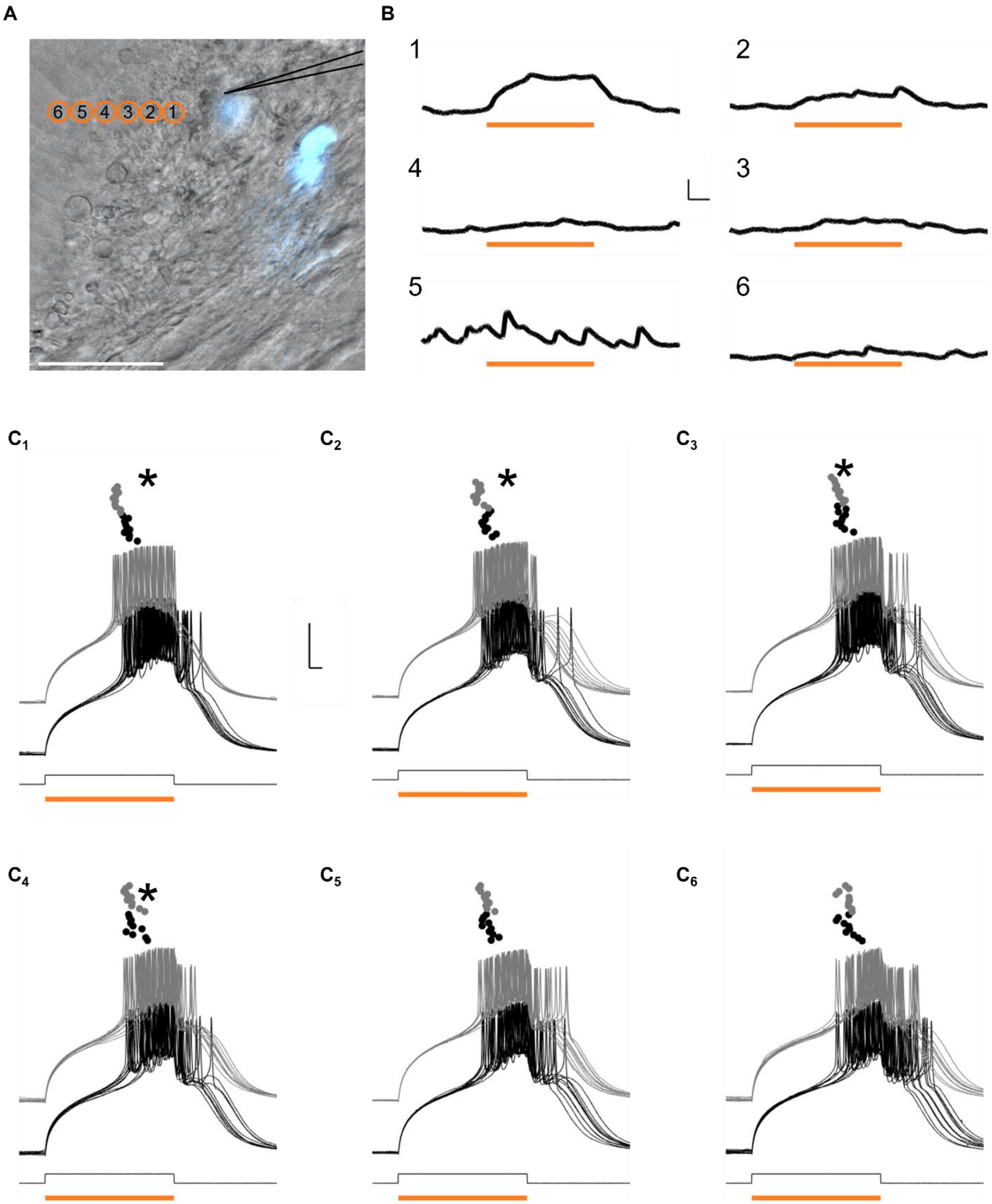


**Fig. S4. Example of excitation from light scattering.**

**A.** Overlay of IR and fluorescence images from live recording of st-ChroME-mRuby-expressing neurons in PV-Cre mouse. Empty locations were serially photostimulated by increasing 10 µm intervals measured from the edge of the soma from the patched neuron. The power of the photostimulation was 10.5 $mW/mm^2$ and was sufficient to drive the neuron to burst when targeted on the soma. Scale bar: 50 μm.

**B.** Voltage responses of the patched neuron from focally stimulating the corresponding numbered locations in A. Scale bar: 10 ms, 1 mV.

**C.** Shifts in in spike latency caused by focally photostimulating empty locations in A. **C1.** $\delta L$ = 13.4%, $p < 0.01$. **C2.** $\delta L$ = 9.2%, $p < 0.01$. **C3.** $\delta L$ = 5.2%, $p < 0.01$. **C4.** $\delta L$ = 3.9%, $p = 0.04$. **C5.** $\delta L$ = 3.5%, $p = 0.07$. **C6.** $\delta L$ = 1.8%, $p = 0.38$. Scale bar: 5 ms, 20 mV. Asterix mark significance ($p < 0.05$, Wilcoxon signed-rank test).

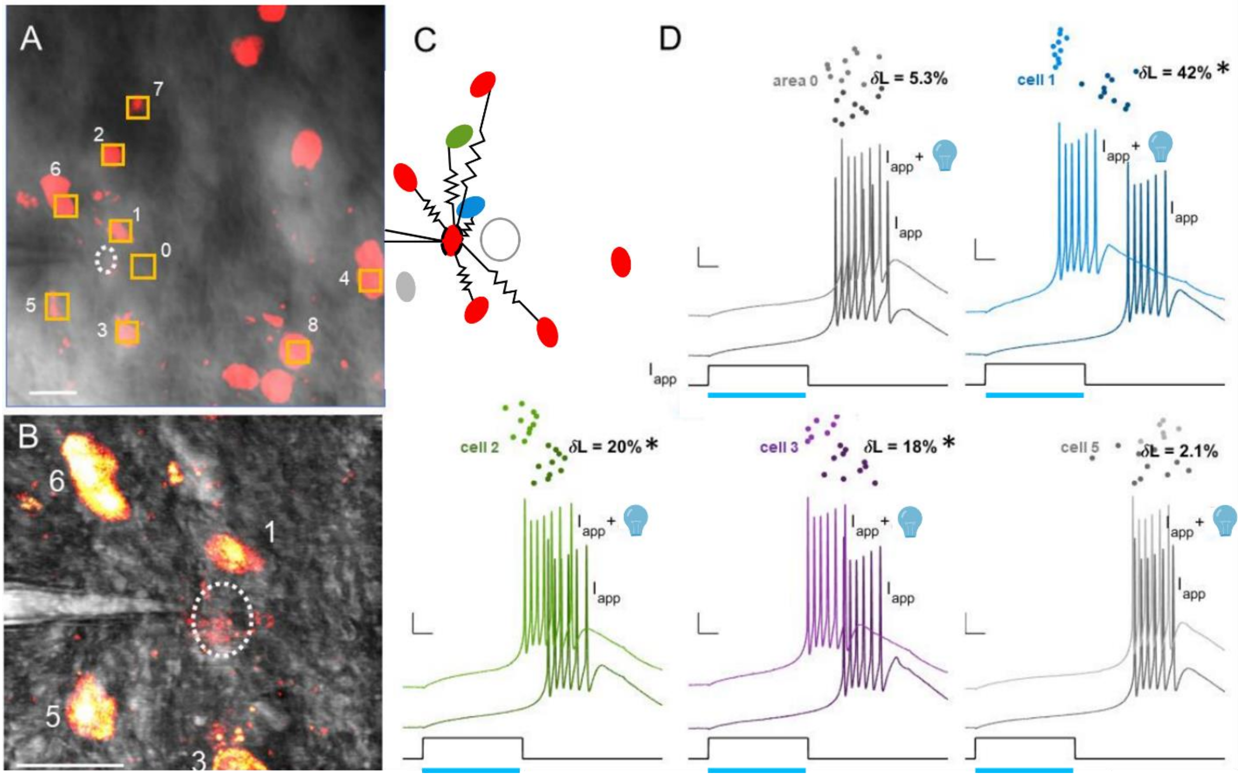


**Fig. S5. Opto-δL mapping of a coupled network with 2-photon stimulation.**

**A.** Composite Dodt + 810 nm z-stack of mRuby2+ neurons in TRN. In this field, one cell was patched (white) and 8 cells and one blank control (0) near it were photostimulated for electrical synapses. 2-photon stimulation was performed via spiral scans (full spirals every 2 ms) ringing the somatic membrane with laser tuned to 990 nm. Scale bar: 20 µm.

**B.** Magnification of recorded neuron and 4 closest cells in A. Scale bar 20 µm.

**C.** Map of coupled network in A as determined opto-δL of areas 0-8 (boxes**).**

**D.** Measurements of δL for areas 0, 1, 2, 3 and 5. Dots represent latency to spike in the patched neuron across 10 trials as driven by a 60-pA current pulse alone (darker colors), and for 10 trial driven by current pulse and simultaneous 960-nm spiral scan stimulation of coupled cell (lighter colors). Scale bars 10 mV, 10 ms. * denotes $p < 0.05$ (unpaired, 2-sided t-test).
